# Heterogeneous timing of asexual cycles in Plasmodium falciparum quantified by extended time-lapse microscopy

**DOI:** 10.1101/344812

**Authors:** Heungwon Park, Shuqiang Huang, Katelyn A. Walzer, Lingchong You, Jen-Tsan Ashley Chi, Nicolas E. Buchler

## Abstract

Malarial fever arises from the synchronous bursting of human red blood cells by the Plasmodium parasite. The released parasites re-infect neighboring red blood cells and undergo another asexual cycle of differentiation and proliferation for 48 hours, before again bursting synchronously. The synchrony of bursting is lost during *in vitro* culturing of the parasite outside the human body, presumably because the asexual cycle is no longer entrained by host-specific circadian cues. Therefore, most *in vitro* malaria studies have relied on the artificial synchronization of the parasite population. However, much remains unknown about the degree of timing heterogeneity of asexual cycles and how artificial synchronization may affect this timing. Here, we combined time-lapse fluorescence microscopy and long-term culturing to follow single cells and directly measure the heterogeneous timing of *in vitro* asexual cycles. We first demonstrate that unsynchronized laboratory cultures are not fully asynchronous and the parasites exhibit a bimodal distribution in their first burst times. We then show that synchronized and unsynchronized cultures had similar asexual cycle periods, which indicates that artificial synchronization does not fundamentally perturb asexual cycle dynamics. Last, we demonstrate that sibling parasites descended from the same schizont exhibited significant variation in asexual cycle period, although smaller than the variation between non-siblings. The additional variance between non-siblings likely arises from the variable environments and/or developmental programs experienced in different host cells.

## Introduction

Malarial fever occurs in multiples of 24 hours. This circadian timing of fevers was already recognized by the ancients even when the cause of malaria remained mysterious. We now know that malarial fever arises from the synchronous bursting of host red blood cells by a eukaryotic parasite (*Plasmodium*). The released parasites called merozoites then infect neighboring red blood cells and undergo another asexual cycle where they use cellular resources to progress through an early ring-stage to trophozoite to schizont for a multiple-of-24 hours before again bursting synchronously. Circadian bursting of parasites from red blood cells is thought to be advantageous because it may help them overwhelm or avoid the host immune response^1,2^. Interestingly, *Plasmodium* species that cause malaria exhibit differences in the timing of their synchronous bursting. For example, the malarial fever that arises from *P. falciparum* has a 48-hour cycle, whereas *P. malariae* has a 72-hour cycle.

However, asexual cycles should lose synchrony with each other as cell-to-cell variability in period leads to increasing differences in the timing and phase of events. This was demonstrated by Trager and colleagues, who revolutionized malaria research by developing a protocol to culture and propagate *Plasmodium* outside of the body^3^, and they showed that the synchrony of bursting was lost during *in vitro* culturing of parasites in flasks. Maintenance of synchronous bursting of *P. falciparum* during malaria infection likely arises from circadian host cues that periodically reset and entrain variable asexual cycles to a similar phase. This interpretation is supported by experiments that inverted the light-dark cycle experienced by hosts and showed that synchronous parasite bursting was subsequently inverted^4–6^. Possible host cues that could affect the asexual cycle include temperature^7, 8^, melatonin^9^ and unknown factors.

Most research uses bulk assays to measure a biological signal averaged over many cells drawn from a population. The lack of synchrony during *in vitro* culturing poses a challenge for studying the asexual cycle because the signal of interest will be averaged over parasites in different stages. Thus, to study the asexual cycle in the laboratory using bulk assays, one must artificially synchronize the parasite population. One common method uses pulses of sorbitol to osmo-shock and kill parasites in the later stages of the asexual cycle and, thus, enrich for ring-stage parasites that span a window of time^10^. This raises the following questions: What is the cell-to-cell variability in the asexual cycle period, and to what extent can it be minimized? Do artificially synchronized parasites exhibit similar cycle dynamics compared to unsynchronized parasites, or does sorbitol shock fundamentally perturb the asexual cycle?

To address these two questions, we employed time-lapse fluorescence microscopy and long-term culturing to image red blood cells infected with a GFP-transgenic parasite for longer than one asexual cycle. We used this single-cell assay to measure the timing and variability in bursting of individual parasites either with or without sorbitol synchronization. By following the bursting of individual parasites and their descendants, we could also quantify whether siblings (merozoites that burst from the same founder schizont) had similar asexual cycle periods to non-siblings (merozoites that burst from different founder schizonts).

## Results

### Tracking parasite bursting using time-lapse microscopy

We modified a recent time-lapse microscopy protocol^11^ to measure the parasite bursting over 80 hours and determine cell-to-cell variability of the asexual cycle in *P. falciparum*. We customized wells in a Corning culture bottle; see Materials & Methods. Briefly, glass slides in the wells were coated with Concanavalin A (0.5 mg/ml), washed with PBS and loaded with 300 *µ*l of red blood cells (RBC) that were infected by synchronized or unsynchronized GFP-transgenic parasites. The synchronized parasites were prepared using two rounds of sorbitol shock^10^. After loading the infected red blood cells, we washed out non-adherent cells, filled the bottle with nutrient medium and blood gas, and left it closed and unperturbed for the remainder of the experiment. We used fluorescence microscopy in a temperature control microscope (37°C) to monitor the timing of burst and re-invasion events, e.g. parasite right before (Fig 1A) and after (Fig 1B) bursting the host RBC. Because parasitemia was low (i.e. few infected RBCs in a field of view) and because merozoites released from a burst locally re-infect RBCs, we could follow a full asexual cycle from the first burst of the schizont “founder” cell (burst time *t*_*i*_ of the i^*th*^ founder) to the subsequent bursts of those RBCs infected by progeny GFP-merozoites (burst time *t*_*i,k*_ of the k^*th*^ progeny descended from the i^*th*^ founder); see Fig 1C, D, E, and F. Such data permit the quantification of variability in the timing of bursts over one asexual cycle; see labeling scheme in Fig 1G. Our observations of “local” infection are consistent with previous work, which found that released merozoites have a limited amount of time to re-infect and, thus, only successfully invade RBCs within one schizont radius^12^.

**Figure 1.**
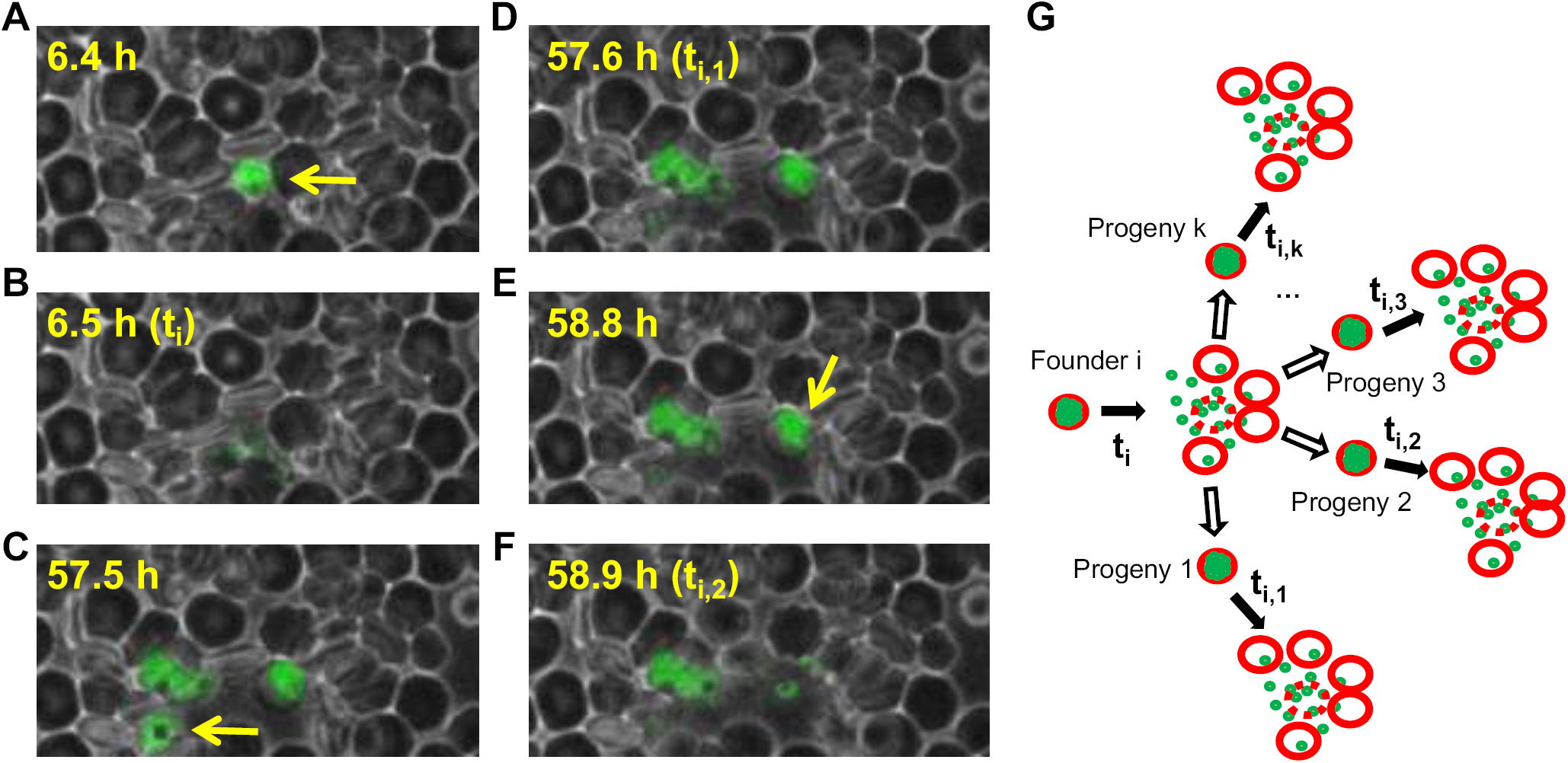
Time-lapse fluorescence microscopy of parasite asexual cycles. (A-F) We measured the asexual cycles of GFP-labeled parasites with phase contrast and GFP fluorescence images taken every 6 minutes (0.1 h). We followed bursting, local reinfection of new red blood cells, asexual growth and division, and re-bursting. (A) Example of an infected red blood cell (e.g. founder cell i) that is about to burst (yellow arrow). (B) Parasite bursting from founder cell i (time of first-burst, *t*_*i*_). (C) Progeny parasites re-infect neighboring red blood cells, proliferate asexually, and the first of the progeny (yellow arrow) is about to burst. (D) Parasite bursting from progeny 1 of founder cell i (time of second-burst, *t*_*i*,1_). (E) The second of the progeny (yellow arrow) is about to burst. (F) Parasite bursting from progeny 2 of founder cell i (time of second-burst, *t*_*i*,2_). (G) Notation for parasite bursting. Founder cell i bursts at time *t*_*i*_, some progeny re-infect red blood cells, undergo an asexual cycle, and burst at time *t*_*i*,1_ (first progeny to burst), *t*_*i*,2_ (second progeny to burst), *t*_*i*,3_ (third progeny to burst), etc. The asexual cycle period (*τ*_*i,k*_) of progeny k of founder cell i is (*t*_*i,k*_ - *t*_*i*_), i.e. burst-to-burst time interval.

### Variability of parasite first-burst times in synchronized and unsynchronized cells

We started by analyzing the elapsed time between the start of the experiment until the first recorded RBC burst of the founder parasites (*t*_*i*_) in synchronized and unsynchronized populations (Fig 2). All experiments were done twice and the first-burst statistics were identical between replicates (Fig S1). In the synchronized culture, 80% of total parasites undergo bursting within the first 10 hours (mean is 6.8 hours and standard deviation is 4.1 hours) displayed either as frequency (Fig 2A) or individual dots (Fig 2B). Surprisingly, the first-burst times of the unsynchronized population were not entirely random or asynchronous (i.e. equally likely to burst at any given time). Rather, the unsynchronized population exhibited two well-defined modes at ∼ 11 hours and ∼ 39 hours. The biological replicate also had a bimodal distribution with similar timing (Fig S1).

**Figure 2.**
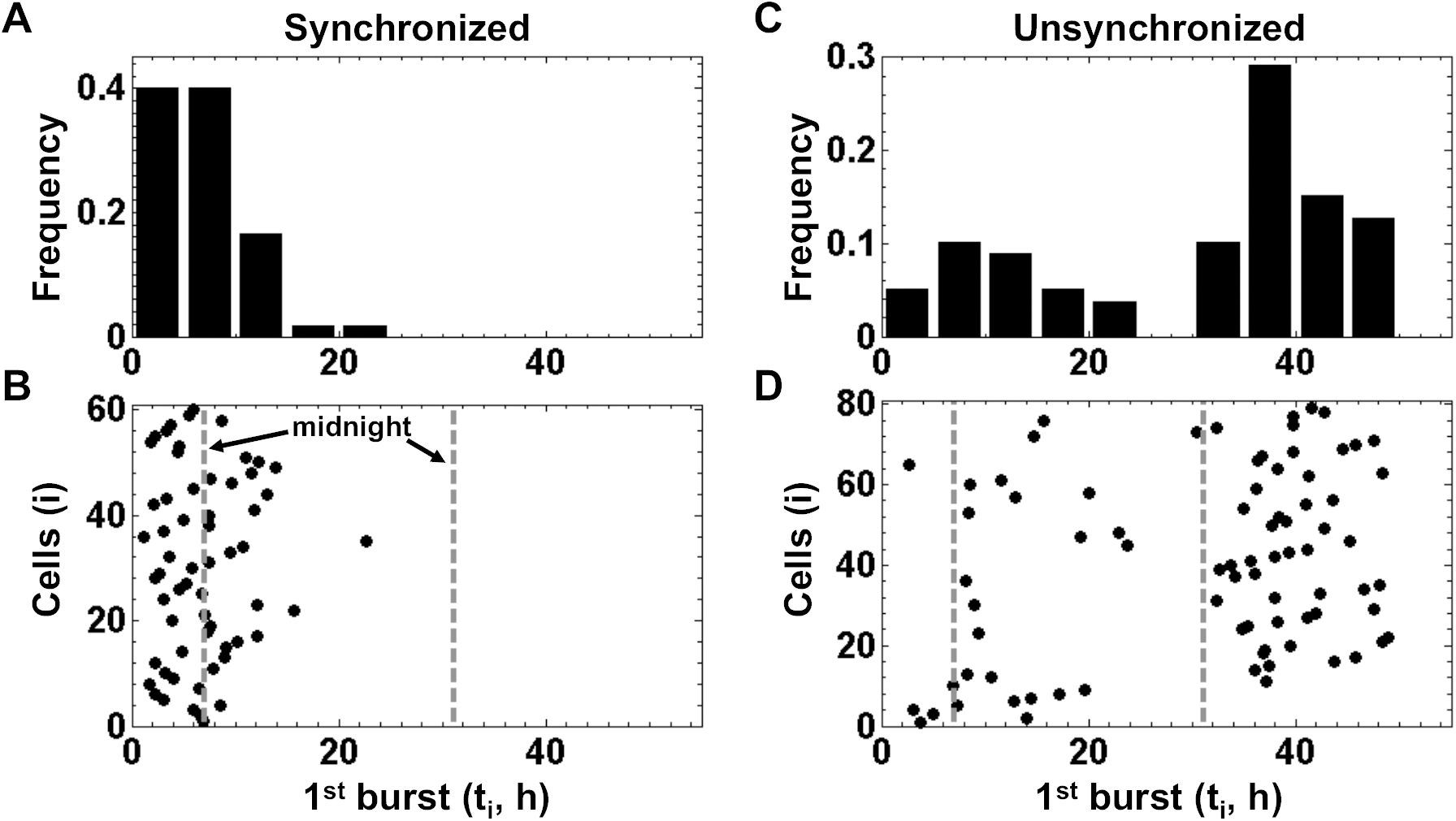
First-burst times of founder cells in synchronized and unsynchronized populations. We plot the first-burst times (*t*_*i*_) of initially-infected founder cells in sorbitol-synchronized (A and B, n=60 founder cells) and unsynchronized (C and D, n=79 founder cells) cultures. B and D plot first-burst times (*t*_*i*_, x-axis) ordered by founder cell (i, y-axis). For circadian reference, perpendicular dotted lines indicate a real world time (i.e. midnight). A and C show the normalized distributions of the burst times in B and D (bin size = 5 hours). Biological replicate is shown in Fig. S1 and complete data are available in Dataset S1. This figure includes founder cells that burst, but whose progeny could not be confidently identified or were unable to re-infect red blood cells (i.e. no *t*_*i,k*_ and, thus, no measured asexual cycle).

### Measuring asexual cycles at the single cell level

Next, we measured asexual cycle periods of parasites with or without synchronization (Fig 3A, C, respectively) by quantifying the difference in the first-burst time of the founder cell (*t*_*i*_, i^*th*^ founder) and the second-burst time of its progeny (*t*_*i,k*_, k^*th*^ descendant of i^*th*^ founder). The asexual cycle periods in the synchronized and unsynchronized parasites were statistically indistinguishable and had a mean of ∼ 55 hours (Fig 3B, D, respectively). The asexual cycle periods of the synchronized and unsynchronized parasites in the biological replicate were similar with a mean of ∼ 58 hours (Fig. S2), which is slightly longer than the first replicate. However, both periods were consistently longer than the synchronous 48-hour asexual cycle of *P. falciparum* that occurs during malarial fever in human hosts, a point that we revisit in our discussion. We also observed that a small fraction of founder RBCs were infected by multiple parasites. We analyzed these multiply-infected RBCs and found that parasites in these multiply infected RBCs have a slightly longer period (55.6 hours) compared to singly-infected RBCs (53.7 hours), although this difference was not statistically significant; see Fig. S3.

**Figure 3.**
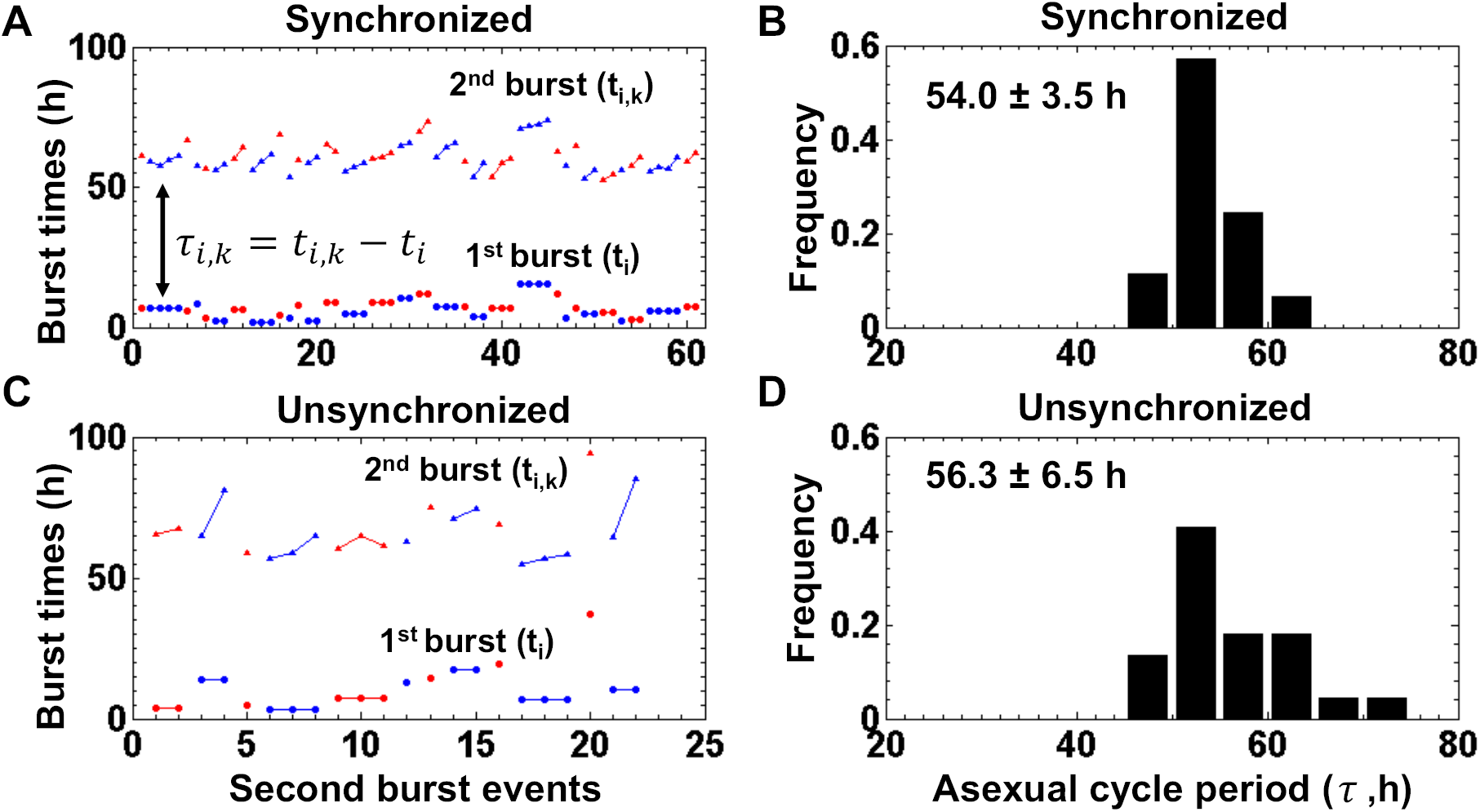
First-burst times of founder cells in synchronized and unsynchronized populations. We plotted the founder first-burst times (*t*_*i*_), progeny second-burst times (*t*_*i,k*_), and asexual cycle period (*t*_*i,k*_-*t*_*i*_) for sorbitol-synchronized (A and B, n=61 founder-progeny pairs) and unsynchronized (C and D, n=22 founder-progeny pairs) cultures. A and C show the first-burst times of the founder cells (*t*_*i*_, dots on y-axis) and second-burst times of the progeny (*t*_*i,k*_, triangles on y-axis) of each founder cell (i, x-axis). Progeny cells from the same founder cell (i.e. siblings) are linked and colored contiguously in alternating patterns of red and blue. B and D show the normalized distributions of asexual cycle period (burst-to-burst interval, *τ*_*i,k*_ = *t*_*i,k*_ – *t*_*i*_) measured from A and C. The numbers show the mean and standard deviation of asexual cycle period. Bin size is 5 hours. Two-sample Kolmogorov-Smirnov test and Wilcoxon rank sum test indicate that asexual cycle period distributions are similar between the synchronized and unsynchronized population (p-value _*KS*_ = 0.39, p-value _*W*_ = 0.31). A biological replicate is shown in Fig. S2, and asexual cycle data are available in Dataset S1.

### Comparison of asexual cycles between siblings and non-siblings

With our unique ability to trace the lineage of the parents and progeny, we sought to compare the asexual cycle periods of siblings (merozoites from the same founder schizont) and non-siblings (merozoites from different founder schizonts). We plotted the asexual cycle time of siblings (grouped by founder) for the synchronized population; see Fig. 4. We found that there were two layers of variation. Siblings, which initially burst from the same founder cell and start tightly synchronized, still exhibit variation in the timing of downstream events and their asexual cycle period. These differences are termed “sibling variance”. In addition, the progeny of different founders exhibit additional variation in their asexual cycle time, which we termed “founder variance”. To estimate these two variances, we considered a simple model *τ*_*i,k*_ = *τ*_*i*_ + *τ*_*k*_ where the first term (asexual cycle time of founder) and second term (sibling variance about the founder asexual cycle time) are drawn from a normal distribution 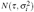 and 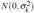, respectively. We first verified that *τ*_*i*_ and *τ*_*k*_ were normally distributed using a Shapiro-Wilks test (p-value > 0.2 for all data sets) and then estimated *τ, σ*_*i*_, *σ*_*k*_; see Table 1. The variability in asexual cycle between siblings (*σ*_*k*_) was less than or equal to the additional variability that arises from founders (*σ*_*i*_) across all conditions and replicates. These data demonstrate that the variability in the asexual cycle period between siblings (*σ*_*k*_) ranges from 2-4 hours and that non-siblings (i.e. most of the population) have an additional source of variability (*σ*_*i*_) due to founder effects.

**Table 1.**
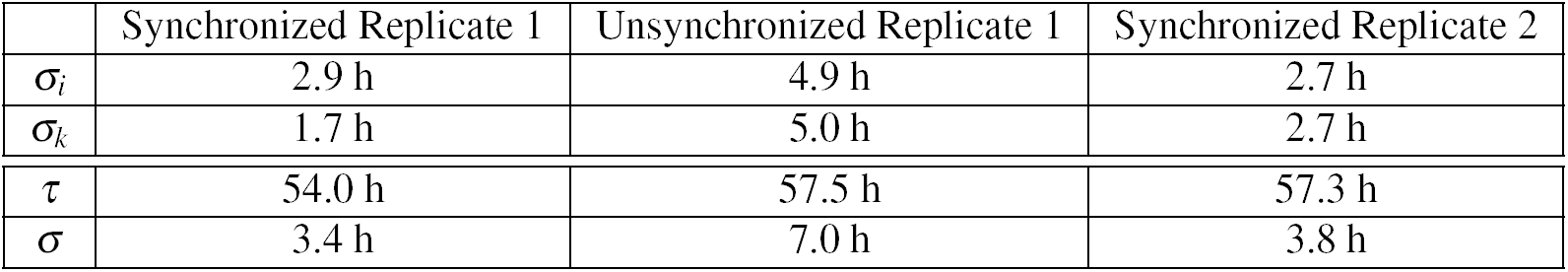
Estimate of founder-specific and sibling-specific variance in asexual cycle period across conditions and replicates. We could not estimate sibling-specific variance in unsynchronized Replicate 2 because it did not have sibling groups (i.e. no doublet or higher). The total variance (*σ*) due to independent contributions from founder-specific and sibling-specific variance is equal to 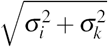.

|  | Synchronized Replicate 1 | Unsynchronized Replicate 1 | Synchronized Replicate 2 |
| --- | --- | --- | --- |
| $\sigma_i$ | 2.9 h | 4.9 h | 2.7 h |
| $\sigma_k$ | 1.7 h | 5.0 h | 2.7 h |
| $\tau$ | 54.0 h | 57.5 h | 57.3 h |
| $\sigma$ | 3.4 h | 7.0 h | 3.8 h |

**Figure 4.**
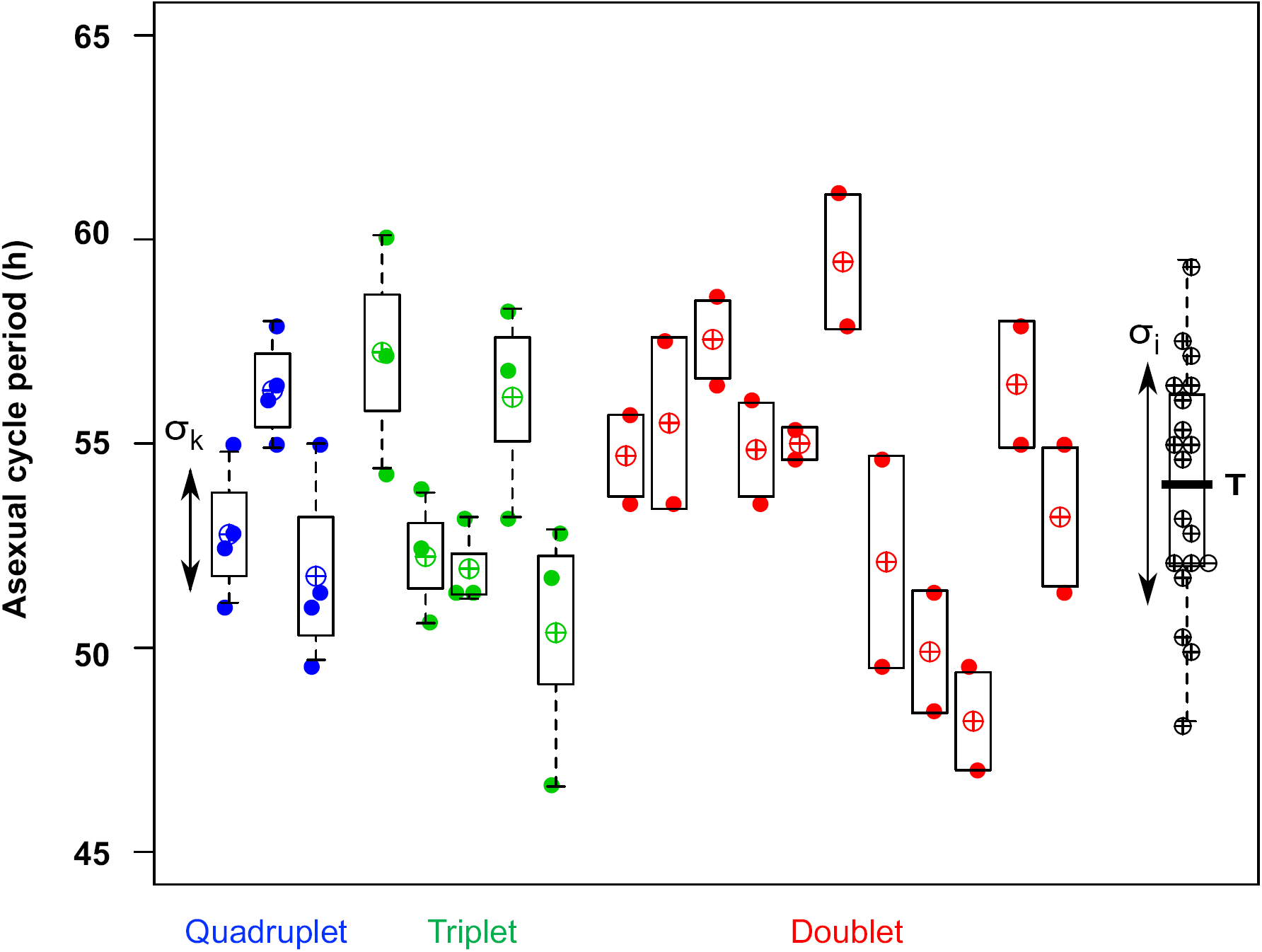
Variance in asexual cycle period between siblings and non-siblings. A plot of asexual cycle period *τ*_*i,k*_ within each sibling group (doublet, triplet, and quadruplet) from the synchronized population in Fig. 3. The mean period of each sibling group (*τ*_*i*_) is shown with circle-plus symbol and projected to the right (black) to display the full distribution of founder times. We estimated the variance in asexual cycle between siblings (*σ*_*k*_) by averaging over all sibling groups. The distribution of *τ*_*i*_ (black) was used to estimate average period (*τ*) and the variance in asexual cycle between founders (*σ*_*i*_).

## Discussion

We used timelapse fluorescence microscopy of GFP-expressing *P. falciparum* to quantify the timing and variability of the asexual cycle in *in vitro* culturing conditions. For the first time, we measured the first-burst time of individual founder parasites, local reinfection, and second-burst times of progeny (Fig. 1) for unsynchronized and sorbitol-synchronized parasites. As expected, the synchronized population exhibited a unimodal distribution of first-burst times with variance due to imperfect synchronization and/or dephasing since the last sorbitol synchronization (Fig. 2A). Surprisingly, the unsynchronized population exhibited a bimodal distribution of first-burst times with two peaks separated by ∼ 28 hours (Fig. 2B and S1). The measured asexual cycle period was unimodal with a mean between 54-58 hours (Fig. 3 and S2), which is longer than the 28-hour difference in the two peaks in the unsynchronized population. The implications are two-fold: (i) *in vitro* cultures may not be perfectly asynchronous, as often assumed, and (ii) our routine culturing appears to create two staggered cohorts of founder parasites in the unsynchronized population. The reason for our bimodal distribution is unknown. One explanation may be that laboratory parasites are cultured by human researchers whose activities are circadian-regulated and our routine lab manipulations synchronize and split the 56-hour asexual cycle population into two cohorts separated by 24 hours. Another possibility is that the parasites self-organize into two separate cohorts via cell-to-cell communication, e.g. exosomes secreted from the infected RBC^13^, such that the split population exhibits a 1:2 relationship with the intrinsic asexual cycle period (28 hour gap between cohorts versus 56 hours asexual cycle). Future experiments could distinguish between these possibilities by culturing parasites in human-free, continuous culture (e.g. bioreactors) and/or changing the mean asexual cycle period by using different strains; see below.

We then quantified the cell-to-cell variability in the asexual cycle period by measuring the time interval between the first burst of each founder and second bursts of its progeny. These single-cell experiments are novel because, for the first time, we measured asexual cycles for parasites in an unsynchronized population. Our results reassuringly show that sorbitol synchronization does not fundamentally perturb the asexual cycle because the mean cycle periods were similar in synchronized and unsynchronized conditions (Fig. 3). The mean period of the GFP-labelled parasites was ∼ 15% longer than 48 hours, a difference that might arise from the burden of GFP expression, RBC immobilization, and/or the photo-toxicity from extended time-lapse fluorescence imaging. To address these concerns, we synchronized and then measured the cycling time of 3D7HT-GFP in standard culture conditions. Our results demonstrate that the asexual cycle period (control) is less than or equal to 49 hours; see Fig. S4. This indicates that the longer cycling time in our time-lapse live fluorescent imaging conditions might be caused by the different culturing and/or imaging condition. To determine whether the light and/or photo-toxicity used in the fluorescent imaging contribute to the longer cycling time^14^, we measured the photon intensity in our microscope and then re-created equivalent photon intensity for parasites in standard culture conditions, where we illuminated culture flasks with a blue LED array every 6 minutes at same photon intensity; see Methods. Our results demonstrate that the photon intensity does not significantly affect the cycling time of 3D7HT-GFP relative to the no light control (Fig. S4). There, the slower cycling time may be due to sub-optimal culture conditions, which is an inherent challenge when measuring multiple cycles of bursting and reinfection over 80+ hours. Possible solutions to this problem might involve the optimization of the culture conditions, including the combination of millifluidic culturing and long-term time-lapse microscopy.

Last, our single-cell assays permitted us to compare sibling and non-sibling asexual cycle times and directly quantify sources of cell-to-cell variability. We made several observations. First, we show that the variability in the asexual cycle period between siblings (*σ*_*k*_) ranged from 2-4 hours (Table 1). This is a fundamental measurement because siblings are as uniformly staged as parasites can possibly get (i.e. same founder cell, same developmental sequence) and this could only be observed by following single parasites. Second, we could show that non-siblings had more cell-to-cell variability in their asexual cycle than siblings because of an additional source of variation specific to each founder (*σ*_*i*_); see Fig. 4 and Table 1. This variation increases the cell-to-cell variation present in a population of parasites, and likely arises from heterogeneities between founders, e.g. age, molecular factors, epigenetic states, or host cell environment. Third, although the mean asexual cycle time was unaffected by culturing conditions, artificial synchronization reduced both sibling and founder variation in the asexual cycle period by two-fold (Table 1). The mechanism for this variance reduction is unknown, but selection for ring-stage parasites could reduce asexual cycle variability by affecting cell physiology and/or the population demographics possibly sensed through cell-cell communication^13^.

We envision extended time-lapse imaging could be applied to answer other questions. For example, one could capture and monitor infrequent, but important, cell-fate events, such as the small proportion (<5%) of parasites that commit to a sexual fate and develop into male or female gametocytes. It would also be possible to use multiple fluorescent and luminescent reporter genes under the control of genes involved in gametocyte commitment to monitor the regulated expression of these genes over extended periods of time or various environmental cues. Extended time-lapse imaging could also be used to measure the wide variety of gene expression and genetic noise inherent in *P. falciparum* and other parasites, as well as the heterogeneous responses of parasites to drug treatment and environmental stress.

## Methods

### Parasite strain and laboratory culturing

Strain 3D7HT-GFP, which constitutively expresses GFP^15^, was obtained from the Malaria Research and Reference Reagent Resource Center (MR4). The reagent was verified by the original contributors before submission to MR4 and we subsequently confirmed phenotype (i.e. GFP expression) with flow cytometry and fluorescence microscopy. Parasites were thawed from frozen stock and cultured in human B+ erythrocytes at 4% hematocrit according to standard procedures^3^ using RPMI 1640 medium with 0.5% AlbuMAX II. Blood was drawn from donors under the IRB protocols approved by the Duke University Health System Institutional Review Board. Research was performed in accordance with relevant guidelines and informed consent was obtained from all participants. Old medium was changed and replaced every 24 hours with fresh RPMI medium, and fresh erythrocytes were usually added every other cycle (48 hours) to keep the parasitemia below 5%. Medium change and/or erythrocyte addition were done in the middle of the day (10 AM - 2 PM) over a period of 1-2 hours in a cell culture hood at room temperature. The cell culture flask was filled with blood gas mixture (5% carbon dioxide, 5% oxygen, balance nitrogen), sealed, and returned to the incubator at 37°C.

### Synchronization of the parasites

To obtain synchronous cultures, parasites were treated with 5% D-sorbitol which selects for parasites in the late ring stage (10). Sorbitol treatment was repeated twice, approximately 48 hours apart, to obtain synchronous cultures. After the last D-sorbitol treatment, synchronized cultures were grown to the trophozoite stage and placed at 5 – 10% parasitemia for imaging as previously described^11^. In brief, 500 *µ*L of parasite culture was centrifuged in a 15-mL Falcon tube at 1000 x g for 30 s on low brake. The supernatant was removed, and the pellet was washed twice in 1 mL of 1x PBS with centrifugations at 1000 x g for 30 s. The pellet was resuspended in 500 *µ*L of RPMI 1640 medium supplemented with 25 mM HEPES, 12 mM NaHCO3, 6 mM D-glucose, 0.5% AlbuMAX II, and 0.2 mM hypoxanthine, and transferred to a 1.5-mL eppendorf tube^16^. In parallel, we prepared unsynchronized cultures from the same initial parasite population.

### Fabrication of the culturing device

We customized a 25 cm^2^ Corning culture bottle for long-term imaging and culturing of samples. We first cut two 10 mm x 10 mm square holes on the bottom of the culture bottle with a spacing of 5 mm. We then sealed a No. 1 glass cover of 25 mm x 40 mm (VWR) to the bottom using aquarium silicone sealant (MARINELAND)^11^ to form two independent wells. After solidifying the sealant overnight, we added 15 mL 70% ethanol for 1 hour to sterilize the surface of wells and interior of the culturing bottle. The ethanol was subsequently removed by vacuuming the device (RobinAir VacuMaster) in a desiccator for 1 hour.

### Time-lapse microscopy

We washed the wells twice using 300 *µ*L 1X PBS (Gibco: DPBS(1x), Dulbecco’s Phosphatase Buffered Saline, [+] Calcium Chloride and [+] Magnesium Chloride). To immobilize the red cells (11) to the glass surface, we coated the well with 0.5 mg/ml Concanavalin A from *Canavalia ensiformis* (Jack bean) Type IV-S (lyophilized powder, aseptically processed, Sigma-Aldrich). Each well was loaded with 300 *µ*L of Concanavalin A solution (dissolved in water) and incubated for 10-20 min at 37°C with the bottle cap fully closed. Concanavalin A was subsequently pipetted out and the wells were washed with 1X PBS twice. We immediately loaded 300 *µ*L of culture synchronized or unsynchronized parasites into each well and incubated for 10 min at 37°C to allow the cells to settle on the surface. Non-immobilized cells were removed and immobilized cells were then further washed by 1X PBS to achieve a monolayer of red cells under the microscope. We added 20 mL of supplemented RPMI 1640 medium. This phenol red-free RPMI medium reduces photo-toxicity^16^. Mixed blood gas was purged into the culture bottle for 1 min at a pressure of about 3 psi before capping and sealing the culture bottle.

We used a temperature-controlled, DeltaVision Elite microscope with an Olympus UPlanFLN M=60x / 1.25 Oil Iris Ph3, Infty / 0.17 / FN=26.5 mm oil objective. Images were acquired with an Evolve EMCCD camera (Photometrics, Tucson, AZ) with 10 MHz speed, 1×1 binning, 10x gain, and 512×512 dpi. Our culturing device was fixed to the microscope stage and the incubation chamber was maintained at 37°C during the entire experiment. To minimize light exposure on the samples, we used 32% transmission efficiency with 0.01-second exposure time for transmitted light, and 2% transmission with 0.025-second exposure for FITC (GFP). Filter set 2 (Polychroic BGmCherry, FR (alexaquad) DAPI, FITC, AF594, CY5) and Alexa mode were chosen to further refine the FITC signal from the microscope settings. Images were collected every 6 minutes for 80 - 120 hours at 40-50 different positions across the two wells. We used the DeltaVision UltimateFocusTM module to compensate for thermal drift and maintain cells in the z-focal plane.

### Image analysis

Phase and GFP images were merged using Image J^17^ to visualize the asexual cycle dynamics of GFP-tagged malaria parasites bursting, reinfecting, proliferating, and re-bursting red blood cells. We subtracted each background from Phase images and GFP images by using the “Subtract Background” function in the Image J and then merged both images in all time-frames. Once we had the time-sequences of the merged images, we checked frame-by-frame to determine the frame in which the parasite burst occurred. During the whole image tracking procedure, we did not change the contrast, but for some frames when parasite bursting was not clear due to low GFP signal, we adjusted the contrast until we got visually clear parasite images and bursting. Our image analysis of asexual cycle dynamics was limited to 80+ hours because uninfected red blood cells started lysing. The spontaneous lysis of red blood cells makes it difficult to distinguish between bursting and lysing in these late time points. We only considered schizont rupture and merozoite release as parasite bursting. Bursting that occurred at earlier stages, such as during the ring or trophozoite stage, was considered hemolysis of the red blood cell.

### Testing the effect of light excitation on the asexual cycle period of GFP-labelled strain

Previous work showed that blue light can perturb the internal pH of malaria parasites^14^. Thus, we performed experiments to test whether GFP excitation light might be responsible for the longer asexual cycle period time observed during time-lapse microscopy. We first measured the total optical power (0.014 mW and 2.43 mW) at the focal plane of our microscope for GFP excitation and phase imaging (T=2% and T=32%, respectively) using a PD300-SH probe attached to an Ophir Nova optical power meter. Our time-lapse samples were illuminated with InsightSSI solid state illumination at 475 nm/14 for 0.025 secs and secs, respectively, every 6 minutes. Thus, the total energy for both snapshots is 2.85 x 10^*-*5^ J or 6.5 x 10^13^ photons with mean photon energy of 475 nm. The 60x phase objective used in our time-lapse experiments illuminated a field of view (FN/M) with a diameter of 442 *µ*m. This corresponds to 4.2 x 10^20^ photons per m^2^ or 0.7 mM photons per m^2^ per snapshot, which is below the photon density known to affect internal pH^14^. However, it is possible that GFP expression, repeated blue light exposure, and the excitation of GFP itself might adversely affect the asexual cycle period of malaria.

To test this, we first grew 3D7HT-GFP in standard culturing conditions and synchronized the strain using the same sorbitol treatment as before, but followed with magnetic sorting when the ratio of rings:schizonts was greater than 2 to enrich for schizonts and further increase the stringency of synchrony. At time zero (t=0 hours), we split the harvested schizonts into two flasks and released them into standard culturing conditions, one of which was illuminated with a blue LED array at the same interval and intensity as used during time-lapse microscopy. We measured the power (17.1 mW per cm^2^) of the blue LED (450 nm) array using a Thors Labs PM200 optical power meter, and determined that a .001 sec exposure every 6 minutes was needed for equivalent photo-illumination conditions. The schizonts were released into dark (control) and light conditions, and we sampled the population every hour between 42 hours and 57 hours to measure new schizonts and subsequent rings expected after one asexual cycle period. Sampled cells were stained with Giemsa, and we counted the total number of parasites in the ring, trophozoite, and schizont stages; see Dataset S2. We fit an error function to data using NonlinearModelFit in Mathematica to estimate the mean and variance in timing of ring emergence; see Fig. S4.

## Acknowledgements

This work was funded by a seed grant from the Duke Center for Genomic & Computational Biology (NEB and JTC), Duke MGM Chair Pilot Fund, Duke Chancellor Award, a National Science Foundation Graduate Research Fellowship (KAW), a National Institutes of Health Director’s New Innovator Award DP2 OD008654-01 (NEB), and a Burroughs Wellcome Fund Pathogenesis (JTC) and CASI Award BWF 1005769.01 (NEB). We thank the Gersbach lab for the use of their blue LED array, the Wax and Kiehart labs for the use of their optical power meters, and Lisa Cameron for help measuring light intensity in the Deltavision Elite microscope located in the Duke LMCF. We also thank an anonymous reviewer for pointing out the multiplicity of infection in Movie S1.

## Author contributions statement

HP, SH, KAW, JTC, NEB conceived of the project; KAW cultured and synchronized strains, performed experiments; HP, SH fabricated culturing device; HP, SH performed time-lapse fluorescence microscopy; HP, NEB analyzed the data; HP, KAW, JTC, NEB drafted initial manuscript; all authors revised and approved of the final manuscript.

## Supplementary materials

**Movie S1**. Timelapse fluorescence movie of GFP-labelled parasites.

**Dataset S1.** First-burst time, second-burst time, and asexual cycle time of synchronized and unsynchronized parasites for both biological replicates. We color-coded MOI=2 (red) and MOI=1 (black) data from Fig. S3.

**Dataset S2.** Ring, trophozoite, schizont counts for a synchronized GFP culture released into control and light conditions.

**Figure S1.**
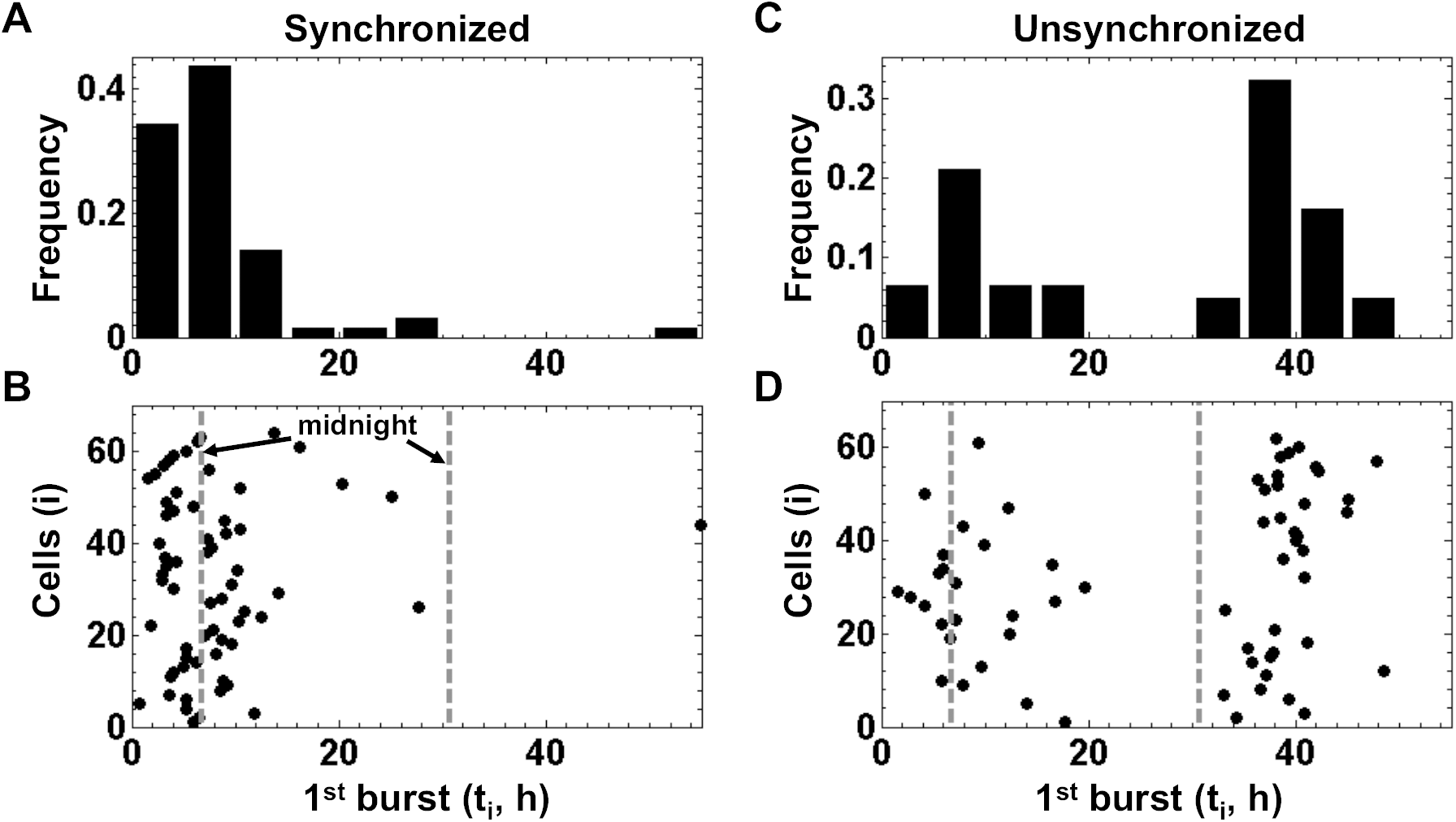
First burst time of founder cells in synchronized and unsynchronized populations are similar between replicates. We plot the first-burst times (*t*_*i*_) of initially-infected founder cells in sorbitol-synchronized (A and B, n=64 founder cells) and unsynchronized (C and D, n=62 founder cells) cultures of a biological replicate. B and D plot first-burst times (*t*_*i*_, x-axis) of founder cells (i, y-axis). For circadian reference, perpendicular dotted lines indicate a real-world time (i.e. midnight). A and C show the normalized distributions of the burst times in B and D (bin size = 5 hours). Biological replicate is shown in Fig. 2, and complete data are available in Dataset S1. Two-sample Kolmogorov-Smirnov test and Wilcoxon rank sum test indicate that synchronized (p-value _*KS*_ = 0.96, p-value _*W*_ = 0.48) and unsynchronized (p-value _*KS*_ = 0.42, p-value _*W*_ = 0.22) first-burst time distributions are similar between replicates.

**Figure S2.**
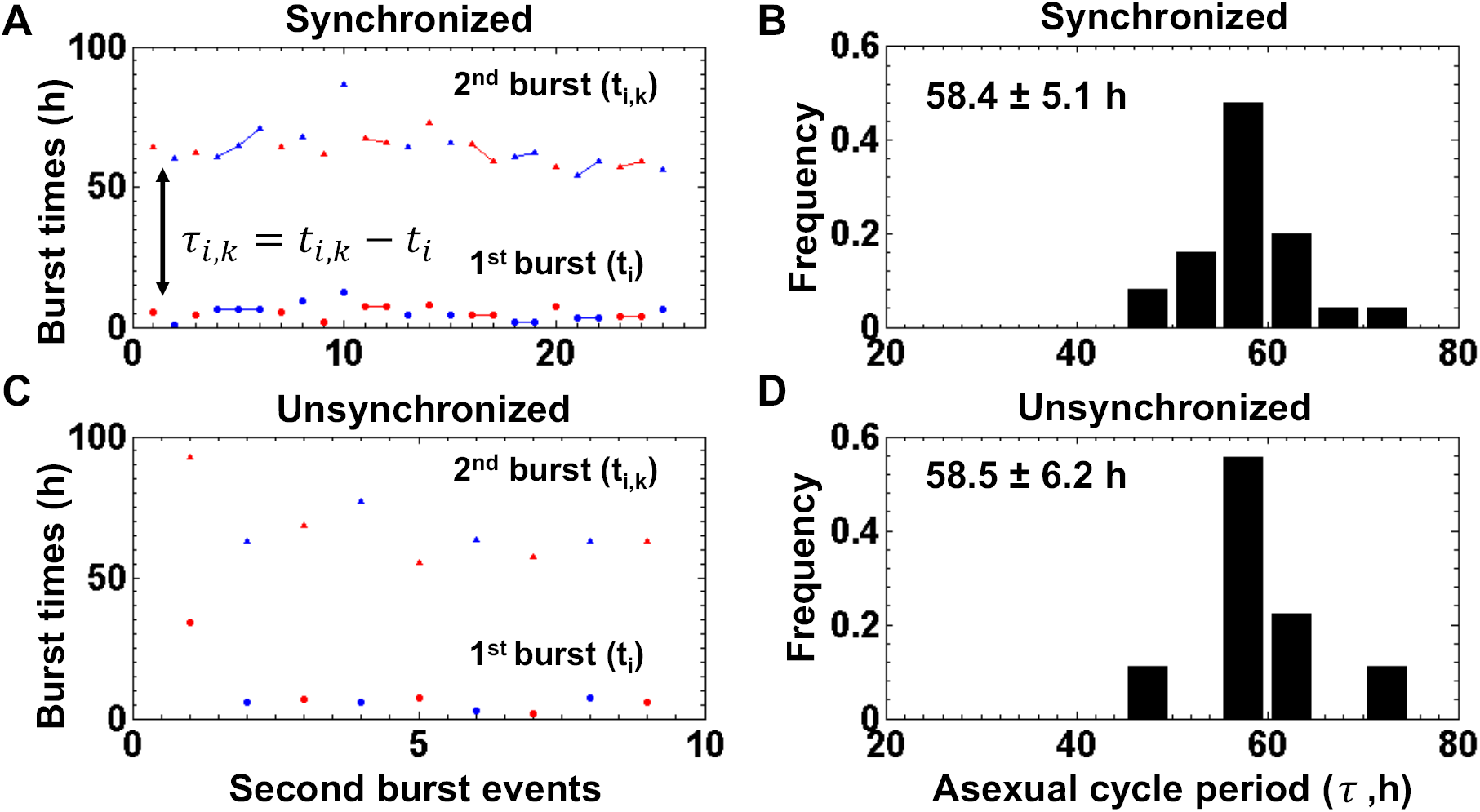
Asexual cycle period in synchronized and unsynchronized populations are similar within a replicate, but differ between replicates. We plotted the founder first-burst times (*t*_*i*_), progeny second-burst times (*t*_*i,k*_), and asexual cycle period (*t*_*i,k*_ - *t*_*i*_) for sorbitol-synchronized (A and B, n=25 founder-progeny pairs) and unsynchronized (C and D, n=9 founder-progeny pairs) cultures. A and C show the first-burst times of the founder cells (*t*_*i*_, dots on y-axis) and second-burst times of the progeny (*t*_*i,k*_, triangles on y-xis) of each founder cell (i, x-axis). Progeny from the same founder cell (i.e. siblings) are linked and colored contiguously in alternating patterns of red and blue. B and D show the normalized distributions of asexual cycle period (burst-to-burst interval, *τ*_*i,k*_ = *t*_*i,k*_ – *t*_*i*_) measured from A and C. The numbers show the mean and standard deviation of asexual cycle period. Bin size is 5 hours. Two-sample Kolmogorov-Smirnov test and Wilcoxon rank sum test indicate that asexual cycle period distributions of the synchronized and unsynchronized population are similar (p-value _*KS*_ = 0.86, p-value _*W*_ = 0.81). However, the distribution of the synchronized population is significantly different between biological replicates (p-value _*KS*_ = 8.6 x 10^*-*5^, p-value _*W*_ = 6.3 x 10^*-*5^). Biological replicate is shown in Fig. 3, and asexual cycle data are available in Dataset S1.

**Figure S3.**
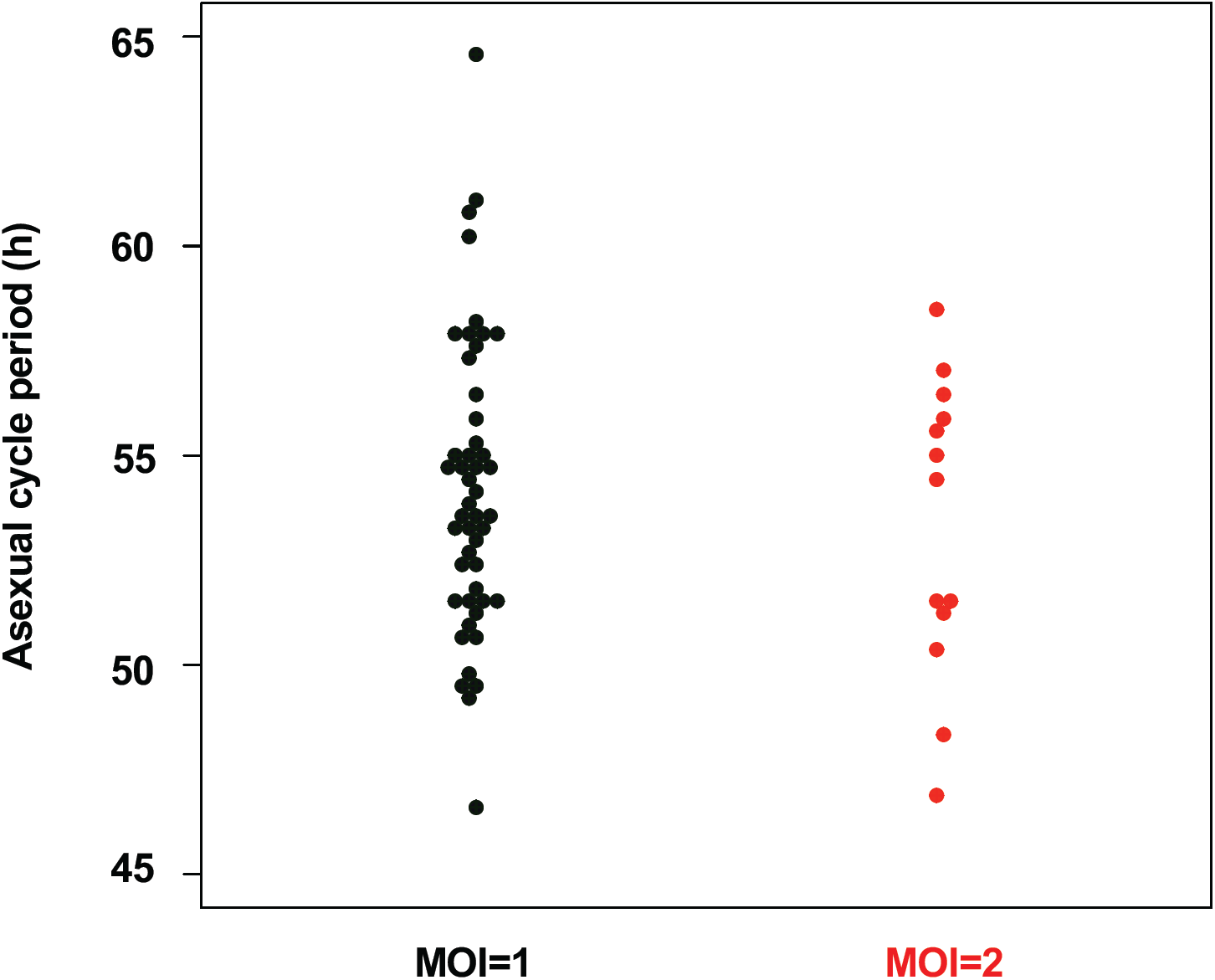
Multiplicity of infection does not significantly affect the asexual cycle period. Some infected RBCs appear to have a multiplicity of infection (MOI) greater than one, i.e. top right corner in Movie S1 at 40 hours. Analysis of the sorbitol-synchronized population from Fig. 3 shows that ∼ 20% (n=13 progeny) have a MOI=2. These MOI=2 progeny appear to have a slightly shorter period (53.0 hours) compared to MOI=1 progeny (54.2 hours), but this difference is not statistically significant. Significance was assessed with two-sample Kolmogorov-Smirnov test and Wilcoxon rank sum test (p-value _*KS*_ = 0.85, p-value _*W*_ = 0.66). Asexual cycle periods calculated from MOI=1 and MOI=2 progeny are indicated in Dataset S1.

**Figure S4.**
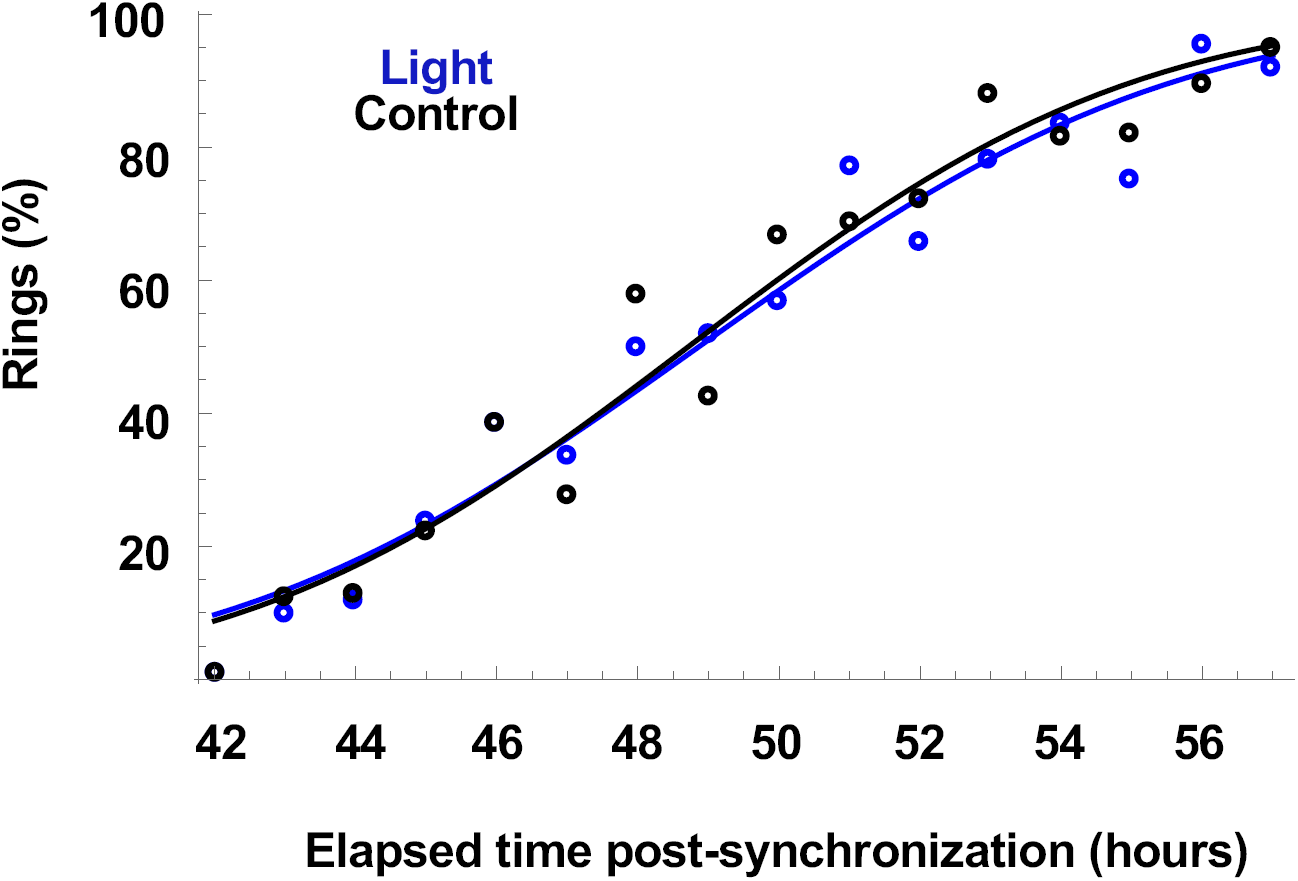
Longer asexual cycle period during microscopy is not due to GFP over-expression and/or photo-toxicity. GFP strains were synchronized by artificially selecting for schizonts using magnetic sorting. These synchronized strains were released into standard culture conditions without light (black circles) and with light (blue circles) at the same intensity and frequency used during our time lapse microscopy; see Methods. We sampled every hour between 42 and 57 hours after release, and counted the number of ring parasites (%); see Dataset S2. Based on Fig. 3, we presumed that timing of the emergence of rings after a full cycle of infection, bursting, and reinfection is well-described by a Gaussian distribution. We then fit an error function (integral of a Gaussian distribution) to the data using nonlinear least squares fitting to estimate mean (*µ*) and variance (*σ*); see Methods. The best-fit curves (solid lines) and parameters for ring emergence (*µ* ±*σ*) are 48.9±5.3 hours (light, blue) and 48.7±5.0 hours (control, black). We conclude that the GFP strain (control) has an asexual cycle period less than or equal to 49 hours, and that blue excitation light does not appear to have an effect on the period. Despite the shorter mean period in standard culturing conditions, the variance is similar to the total variance measured by time-lapse microscopy; see Table S1.

